# Universal chromatin state annotation of the mouse genome

**DOI:** 10.1101/2022.12.19.521116

**Authors:** Ha Vu, Jason Ernst

## Abstract

Genome-wide chromatin states learned from integrating genome-wide maps of multiple epigenetic marks within the same cell type have been widely used to generate genome annotations of individual cell types. An alternative strategy based on ‘stacked modeling’ can provide a single ‘universal’ chromatin state annotation based jointly on data from many cell types. In human, such an approach was recently demonstrated and the resulting chromatin state annotation, denoted full-stack, was shown to have complementary advantages to per-cell-type annotations. However, an analogous annotation has not been previously available in mouse. Here, we produce a chromatin state annotation for mouse based on 901 datasets assaying 14 chromatin marks in 26 different cell or tissue types. To characterize each chromatin state, we relate the states to other external annotations and compare them to analogously defined states in human. We expect the full-stack chromatin state annotation for mouse will be a useful resource for studying the genome of this key mammalian model organism.

## Background

Mouse is widely adopted as a model organism for human for many reasons including their genetic and physiological proximity to humans, relatively short life span, and availability as test subjects for genetic manipulations [1–3]. A wealth of epigenomic datasets in mouse, include maps of histone modifications and variants and sites of accessible DNA, has accumulated thanks to efforts from different consortia and individual labs, which can be used to annotate the mouse genome, including non-coding regions [4–10]. This type of data has previously been integrated methods such as ChromHMM and Segway [11–14] to generate chromatin state maps for various organisms including different mouse and human cell and tissue types [6,15–19]. These chromatin state maps have traditionally been used to annotate genomes in a per-cell-type manner using either the ‘independent’ or ‘concatenated’ modeling approaches (for ease of presentation, we will refer to tissue types also as cell types) [14,20].

Recently, we applied an alternative ‘stacked’ modelling approach of ChromHMM to learn chromatin states from over 1000 human datasets representing more than 100 cell types, to generate a universal annotation of the human genome that can annotate all human cell types [21]. This modeling provided a single annotation of the genome per position based on data from all the input cell types. Such an annotation, denoted full-stack annotation, offers complementary advantages to per-cell-type annotations, such as differentiating constitutively active regions from cell-type-specific ones and simplifying genome annotations across cell types through a single annotation shared across cell types as opposed to one for each. Additionally, the full-stack annotation allows researchers to bypass picking a single cell type for analyses or conducting analyses separately for every cell type. This can be particularly useful in studies involving data that is not inherently cell-type-specific such as analyses of genetic variants or conserved DNA sequence. However, an analogous full-stack annotation has not been previously available in mouse.

To address this, we train a full-stacked model with ChromHMM using input data from >900 mouse datasets of 14 chromatin marks from 26 mouse cell type groups (**Methods**). We analyze these states with respect to their enrichments with external datasets and annotations to provide detailed characterizations for each state. We also analyze to what extent each state is conserved in human. We expect the mouse full-stack annotations along with the provided biological characterizations will be a useful resource for studying this key model organism.

## Results and discussion

We learned the mouse full-stack model by applying ChromHMM to over 900 mouse epigenomic datasets, similar to how it was previously applied in human [12,21] (**Methods, Fig. 1, Fig. S1**). We used a 100-state model for consistency with the previously analyzed human full-stack model. We manually grouped these 100 states into 16 groups. One of the groups contains states associated with assembly gaps or alignment artifacts (mGapArtf), the latter of which are often marked by signals of both open-chromatin mark (ATAC or DNase) and heterochromatin mark H3K9me3 (**Fig. 1**). Another group, Quiescent group (mQuies), consists of states associated with minimal signals of any chromatin marks. We defined a Heterochromatin (mHET) group primarily associated with H3K9me3, and a Zinc finger genes (mZNF) group associated with both H3K36me3 and H3K9me3. We also defined a Polycomb repressed group (mReprPC) associated with primarily H3K27me3, and another group associated with both open chromatin marks (DNase and/or ATAC-seq) and polycomb-repressed-associated mark H3K27me3 (mReprPC_openC). We also defined a group of states associated with just open chromatin (mOpenC), based on DNase-seq and ATAC-seq signals relative to other chromatin marks.

**Figure 1:**
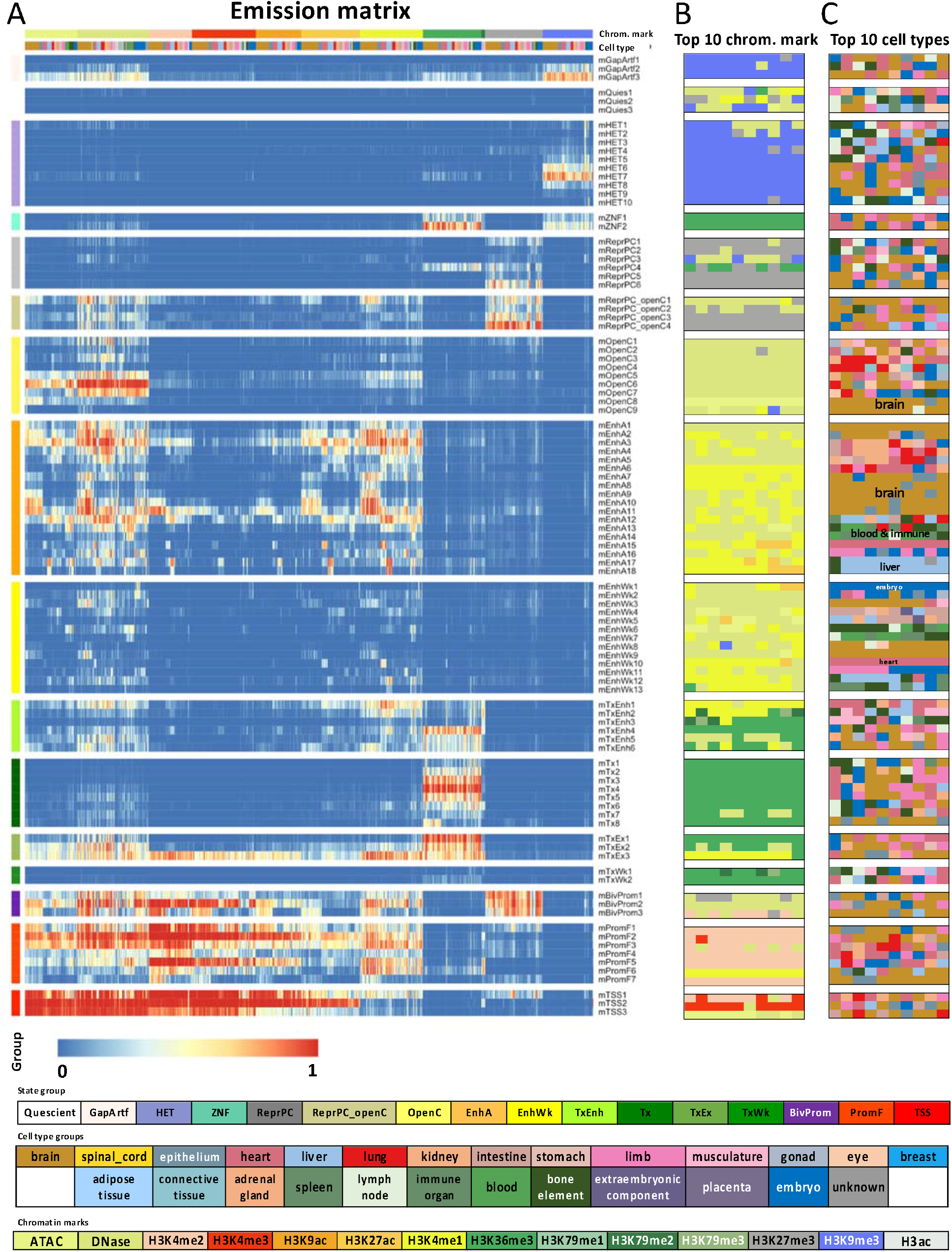
Mouse full-stack state emission parameters. **(A)** Each of the 100 rows in the heatmap corresponds to a mouse full-stack state. Each of the 901 columns corresponds to one input dataset. For each state and each dataset, the heatmap gives the probability within the state of observing a binary present call for the dataset’s signal. Above the heatmap, one color bar indicates the assay/chromatin mark measured by each dataset. The other color bar shows the cell type groups associated with each dataset. The corresponding color legends are shown towards the bottom. The states are displayed in 16 groups with white space between each group, and grouped based on biological interpretations indicated by the color legend at the bottom. Full characterization of states is available in **Additional File 2.** The model’s transition parameters between states can be found in **Additional File 1: Fig. S1.** Columns are ordered such that datasets profiling the same chromatin marks are next to each other. **(B)** Each row corresponds to a full-stack state as ordered in (**A**). The columns correspond to the top 10 datasets with the highest emission value for each state, in order of decreasing ranks, colored by their associated chromatin marks as in (**A**). **(C)** Similar to **(B)**, but datasets are colored by the associated cell type groups. The tissue groups primarily associated with some of the enhancer states are noted inside the heatmap.

We defined three groups of states associated with enhancers: active enhancers (mEnhA), weak enhancers (mEnhWk), and transcribed enhancers (mTxEnh). States in the mEnhA group were associated with open chromatin, H3K27ac and H3K4me1. States in the mEnhWk group (mEnhWk) also showed association with those marks, but at lower levels compared to those in mEnhA group. States in mTxEnh group showed signals of open chromatin (ATAC and/or DNase), H3K4me1, H3K27ac *and* transcription-associated marks (H3K36me3 or H3K79me2/3).

In addition to mTxEnh group, we defined three additional transcription groups: transcription (mTx), transcription and exons (mTxEx), and weak transcription (mTxWk). States in the mTx group are associated primarily with the transcription marks H3K36me3 and/or H3K79me2/3. Meanwhile, states in transcription and exon group (mTxEx) are associated with both open chromatin and transcription marks. States in the mTxWk group are associated with low levels of the transcription marks.

We also defined three promoter-associated groups: bivalent promoters (mBivProm), promoter flanking (mPromF) and transcription start sites (mTSS). States in these groups generally had relatively high levels of H3K4me2 and H3K4me3, and for some of them also H3K4me1 and/or open chromatin marks. mBivProm states were also associated with the repressive mark H3K27me3. States in the mTSS group tended to have weaker H3K4me1 levels.

Within each group, there were differences among individual states, such as the magnitude of the emission probabilities associated with specific chromatin marks, or their association with different cell type groups (**Fig. 1**). For example, different states in the active (mEnhA) and weak enhancer (mEnhWk) groups have enhancer associated marks that were specific to different cell type groups such as the brain, blood, immune, liver, and embryo (**Fig. 1C**). Detailed descriptions of each state’s chromatin mark signals and cell-type-specific activities are provided in **Additional file 2**.

We also conducted various enrichment analyses to further characterize the states (**Fig. 2A**). Enrichments with external annotations further highlight the distinctions among states from different groups, as well as among those within the same group. For example, the state mGapArtf1 overlapped with 99.9% of annotated assembly gaps in mm10 (6.6-fold) (**Fig. 2A**). States mGapArtf1 and mGapArtf3 jointly overlapped with 82.1% of the blacklisted regions from ENCODE (5.4 and 5.0-fold, respectively) (**Fig. 2A**). States in promoter-associated groups (mTSS, mPromF, mBivProm) showed relatively high enrichments with regions within 2kb of annotated TSSs (9.4-26.7 fold, **Fig. 2A**). These states vary in their enrichments with regions upstream and downstream of annotated TSSs (**Fig. 2D, Additional File 1: Fig. S2**). Three states from the TSS group (mTSS1-3) had the strongest enrichment for TSS (59.2-159.9 fold). These three states along with mBivProm2 were strongly enriched with CpG Islands (101.1-159.2 folds, **Fig. 2A**). States in the transcription associated groups (mTx, mTxWk, mTxEnh, mTxEx) all had enrichments greater than 2.4-fold for annotated gene bodies. States in the transcription and exon group (mTxEx1-3) showed the highest enrichments for annotated exons (11.3-14.7 folds, **Fig. 2A**) and regions surrounding annotated TESs (**Fig. 2E, Additional File 1: Fig. S2**). States mOpenC6-7, which had strong *constitutive* DNase-seq and/or ATAC-seq signal while having relatively limited histone modification signals, had the strongest enrichments with CTCF binding sites in multiple cell types (geometric mean 146- and 98-fold for states mOpenC6-7, respectively) (**Fig. 1, 2F, Additional File 3**).

**Figure 2:**
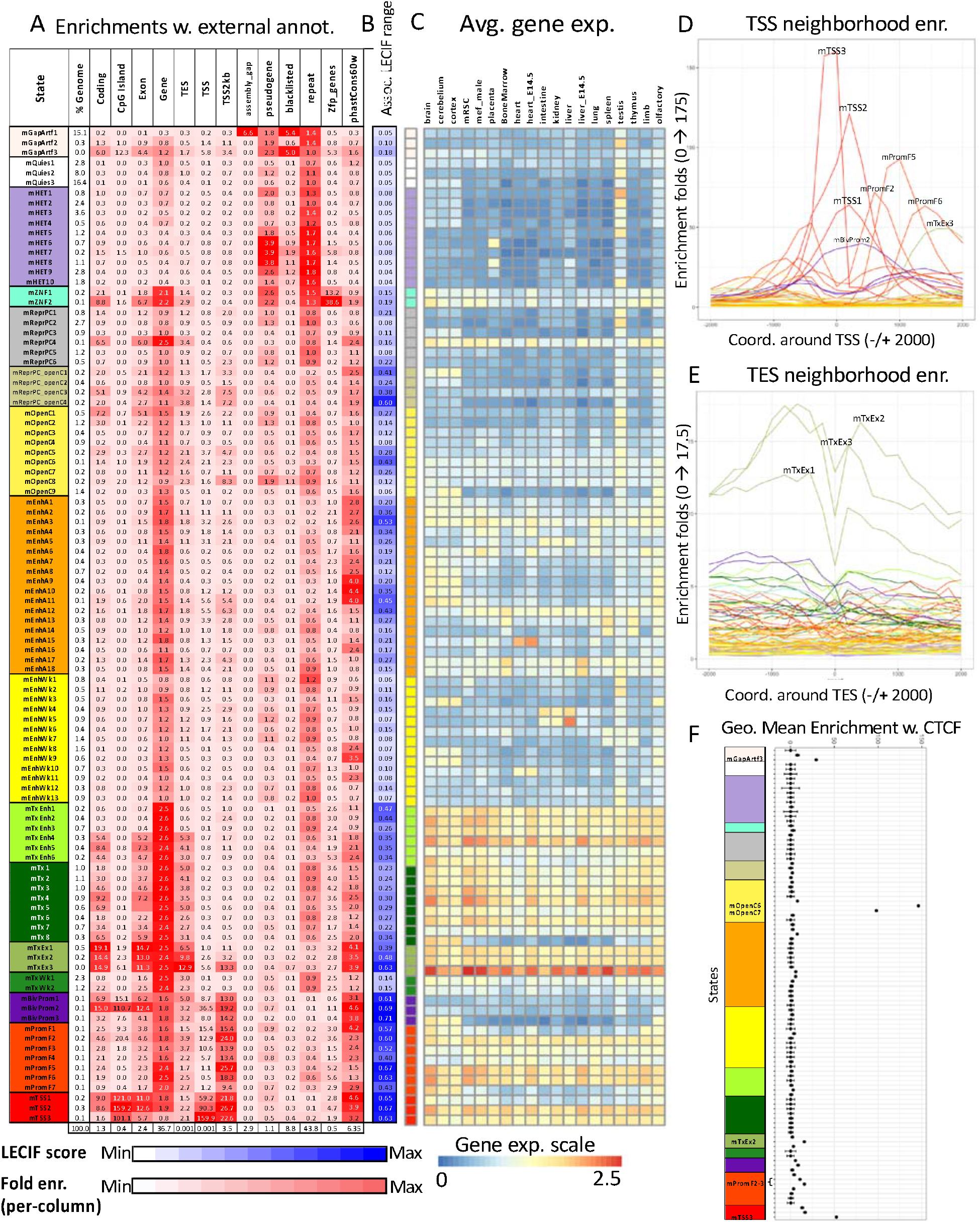
Mouse full-stack states enrichments for external genomic annotations. **(A)** Fold enrichments of mouse full-stack states with external genome annotations (**Methods**). Each row corresponds to a state and each column corresponds to one external genomic annotation: coding sequences, CpG Islands, Exons, gene bodies (exons and introns), transcription end sites (TES), transcription start sites (TSS), TSS and 2kb surrounding regions, assembly gaps, pseudogenes, blacklisted regions, repeat elements, annotated Zpf genes and PhastCons conserved elements (**Methods**). The last row shows the percentage of the genome that each external genome annotation covers. The heatmap colors are column-normalized, i.e. within each column, the colors of the cells are such that highest values are colored red and lowest values are colored white. **(B)** Each row indicates the states’ average LECIF score, indicating functional human-mouse conservation based on epigenetic annotations (Kwon and Ernst, 2021) (**Methods**). The list of states with top average LECIF scores and highest enrichments with PhastCons elements is in **Additional File 1, Fig. S9** and **Additional File 4**. **(C)** Average weighted expression of genes that overlap each full-stack state in different groups of cells (**Methods**). Each column in the heatmap corresponds to a cell group indicated at the top. Each row corresponds to a state, as ordered in **(A).** **(D-E)** Positional enrichments of full-stack states relative to annotated **(D)** transcription start sites (TSS) and **(E)** transcription end sites (TES). Positive coordinate values represent the number of bases downstream in the 5’ to 3’ direction of transcription, while negative values represent the number of bases upstream. Each line shows the positional enrichments in a state. Lines are colored corresponding to the state group as indicated in **(A**). **(F)** Geometric mean and geometric standard deviation of enrichments of full-stacks states CTCF elements across 28 cell types from ENCODE [15] (**Methods**). States are displayed vertically in the same order as **(A).** The DNase6-7 state showed the strongest enrichment for CTCF elements in all observed cell types. The geometric mean and standard deviation are calculated such that for each state, fold enrichment values of 0 are replaced by the state’s minimum non-zero value. The fold enrichment values accompanying this plot are available in **Additional File 4.**

Additionally, we analyzed the enrichment of full-stacked states for different chromosomes. This uncovered three states in the polycomb repressed group (mReprPC4-6) that were highly enriched on chromosome X (8.9-11.4 fold, **Additional File 1: Fig. S3**), likely related to H3K27me3-associated chromosome X inactivation [22,23]. We also found chromosome Y strongly enriched for mGapArtf1 state (6.4 fold, corresponding to 96% of chrY) (**Additional File 1: Fig. S3**).

We also analyzed the states’ enrichments for different classes of repeat elements [24]. For the two largest classes of repeats, Long interspersed nuclear elements (LINE) and long tandem repeats (LTRs) (**Additional File 1: Fig. S4-5**), the most enriched states were both in the HET group (mHET9,7) (2.7 and 3.3 fold). Satellite and rRNA had the strongest enrichments for the mGapArtf3 state, 22.5 and 95.5 fold, respectively.

We also related the full-stack states to average expression of overlapping genes (**Methods**). States in the transcription-associated groups (mTxEnh, mTx, mTxEx), along with those related to promoter (mPromF and mTSS groups) showed higher average gene expression across cell types compared to other groups (**Fig. 2C**). State mTxEx3 showed the highest gene expression of all states.

Additionally, we analyzed the mouse full-stack states’ association with per-cell-type chromatin state annotations defined across 66 reference epigenomes from 12 unique cell type groups and 7 developmental stages, based on 8 marks (**Additional File 1: Fig. S6-7, Additional File 5**) (Gorkin et al., 2020). This revealed, for example, that state mEnhA17 showed the strongest enrichments with per-cell-type active enhancer states across all developmental stages for liver (**Additional File 1: Fig. S6-7, Additional File 5**), which is consistent with this state’s highest signals in enhancer-associated chromatin marks (H3K4me1, H3K27ac) for liver datasets (**Fig. 1**). State mTSS2 was most enriched with per-cell-type active promoter states in all reference epigenomes (**Additional File 1: Fig. S6-7, Additional File 5**), consistent with its association with individual chromatin marks (**Fig. 1**).

In addition, we analyzed how the mouse full-stack states correspond to those of an analogous previously defined full-stack model in human [21]. We evaluated the enrichments of each mouse full-stack state with each human full-stack state after mapping the human annotations to those for mouse (**Methods, Additional File 1: Fig. S8-9, Additional File 4**). Twenty-two out of 100 mouse states showed >50-fold enrichment with at least one human state (**Additional File 1: Fig. S9, Methods**) [21], and these states’ biological implications highlight strong correspondence of states from the human and mouse models. For example, mouse state mTxEx3 showed 378.8-fold enrichment for human state TxEx4 – the largest enrichment across any pair of states—(**Additional File 1: Fig. S9, Additional File 4**). These two states showed the highest average gene expression across multiple mouse and human cell types, respectively [21] (**Fig. 2C**). All 13 mouse states in the promoter groups (mPromF, mBivProm, mTSS states) showed strong enrichments with human full-stack states that are also promoter-associated, with 12 of these mouse states showing >90-fold enrichment (**Additional File 1: Fig. S9, Additional File 3**). Mouse states mOpenC6-7, which are associated with *constitutive* open chromatin CTCF elements (**Fig. 2F, Additional File 3**), showed the strongest association with the human DNase state, which was also constitutively marked by DNase and CTCF in human [21]. However, there exist differences between states from the two organisms’ models. For example, in the mouse model, seven states in the mOpenC group (all except mDNase6-7), which we characterized as showing cell-type-specific signals of open chromatin, did not show strong enrichment for specific human states (**Additional File 1: Fig. S8-9, Additional File 4**).

We also evaluated each full-stack state’s average human-mouse LECIF score, which quantifies conservation at the functional genomics level between the two species (**Fig. 2B**) [25] (**Methods**), which ranged from 0.04 (mHET9) to 0.71 (mBivProm3) (**Fig. 2B, Additional File 1: Fig. S9**). All 14 mouse states that had an average LECIF score >= 0.5 also had a >50-fold enrichment with a human full-stack state, highlighting that mouse states with high LECIF score show concordance with specific human states. In addition, we looked at each state’s enrichment for sequence constraint eleements as defined by PhastCons [26]. Across all states, the states’ enrichments for PhastCons elements and average LECIF score showed overall consistency (Spearman correlation 0.70; p-value: 3.8e-16). We found 10 mouse states that are among the top 20 states based on average LECIF score, enrichments for PhastCons element and for a specific human full-stack state (**Additional File 1, Fig. S9**). Among these states, seven are associated with promoter activities (mBivProm1-3, mTSS1-3, mPromF1), two states are characterized by strong exon enrichments and constitutive transcriptional activities (mTxEx2-3), and one state (mEnhA3) corresponds to constitutively strong enhancers (**Additional File 1: Fig. S9**). Interestingly, a few states stand out as associated with either high sequence constraint or functional conservation (LECIF score), but not in both. For example, constitutive DNase-candidate insulator states mOpenC6-7 are among top 20 with highest average LECIF scores yet had lower (Phastcons) sequence constraint enrichment (ranked 50, 59) (**Additional File 1: Fig. S9**).

## Conclusions

We introduced the mouse full-stacked annotation to provide a single chromatin state annotation per genomic position based on over 900 epigenomic datasets representing 26 different cell type groups. The mouse full-stacked model and its characterization is analogous to the previous human full-stack model [21] (**Data Availability, Additional File 2**). As discussed previously in the context of the human genome annotation [21], the full-stack model has a number of advantages, such as being able to differentiate constitutive from cell type-specific annotations and simplifying the overall genome annotation in that there is a single genome annotation per position. However, this does come at a trade-off of a more complex set of model parameters. The full-stack annotation is not meant to replace existing per-cell-type annotations, but rather to complement them and the most appropriate annotation will likely depend on the application [21]. We expect the full-stack model to serve as an additional resource for work that leverages the mouse as a model organism to gain insight into human biology and disease.

## Methods

### Input data and processing

We obtained data of ENCODE Project Portal [5,6,27], and restricted the downloaded files to those with ‘File analysis title’ starting with ‘ENCODE4’ and ‘File assembly’ of ‘mm10’. In total, we downloaded data of read alignment (.bam files) for 901 experiments, 114 of which were DNase-seq, 83 were ATAC-seq and 704 were ChIP-seq data targeting 12 chromatin marks representing 26 cell type groups (**Additional File 2**). For each .bam file resulting from a ChIP-seq assay, we extracted the corresponding control .bam file by matching the .fastq files of reads from the ChIP-seq assay with the control reads. As the DNase-seq or ATAC-seq experiments did not have paired control .bam files, we assumed a uniform background read distribution. Links to download all input data for the stacked model are provided in **Additional File 2**.

We then constructed the cell_mark_file input table required by ChromHMM BinarizeBam such that there are four tab-delimited columns in the table. The first column is set as ‘Genome’ across all rows. The second column denotes the experiment names of the form ‘<Biosample term name>_<Experiment target>_<Experiment accession>’, where ‘Biosample term name’, ‘Experiment target’ and ‘Experiment accession’ correspond to the cell type, histone mark/DNase/ATAC profiled and the accession code of such experiments, respectively from the metadata from ENCODE. The third column contains the experiments’ .bam file names. The last column contains the matched control .bam file names, which is left blank for DNase-seq or ATAC-seq experiments, since we assumed a uniform background distribution for these assays. Using this cell_mark_file input table, we next binarized the data at 200 base pair resolution using the BinarizeBam and MergeBinary commands of ChromHMM (v.1.23), following the procedures of [21].

### Training full-stack modeling and generating genome-wide state annotations

We learned the mouse full-stack chromatin state model for the 901 datasets using the LearnModel command of ChromHMM (v.1.23). We applied the same set of flags as in learning the human full-stack model (-splitrows -holdcolumnorder -pseudo -many -p 6 -n 300 -d -1 - lowmem -gzip), described in Vu and Ernst, 2022 [21]. We specified the number of states to be the same as in the human model (100 states) [21].

### Enrichment and estimated probabilities of overlap with per-cell-type chromatin state annotations

We obtained per-cell-type 15-chromatin state annotations for 66 reference epigenomes/cell types from Gorkin et al., 2020 [19], with download links provided in Kwon and Ernst, 2021 [25]. For simplicity, we use reference epigenome and cell types interchangeably, and we refer to the chromatin state segmentation that is used to annotate the individual reference epigenomes as per-cell-type annotations. This model was trained using the *concatenated* modeling approach from data of 8 chromatin marks measured in 12 cell type groups at up to 8 distinct stages during mouse fetal development [19]. We applied the same procedure as outlined in Vu and Ernst, 2022 [21] to obtain two types of summary results of the relationship between mouse full-stack states’ association with states in per-cell-type annotations. First, for each full-stack state, we report, for each of the 64 reference epigenomes, the chromatin state from the per-cell-type model that is maximally enriched in the full-stack state [21]. Second, for each of the 12 tissue types, we report the estimated probabilities of each full-stack state overlapping with each of the 15 states in the per-cell-type model [21]. These results, along with detailed comments about the observed patterns of overlap between each full-stack state and per-cell-type state, are available in **Additional File 5.** Data of all per-cell-type annotations are in mm10 [19].

### Average gene expression associated with each full-stack state

We obtained data of gene expression for 19 tissue types in mouse from [28] (http://chromosome.sdsc.edu/mouse/download/19-tissues-expr.zip). The provided data contains two gene expression datasets for each tissue type, corresponding to two replicates. We converted the gene expression values for the 19 tissues into *log*_2_ (*FPKM* + 1) values, where FPKM (Fragments per kilo base of transcript per million mapped fragments) were the provided values from the source data, and we added a pseudo count of 1 for each value.

Since the gene expression data was provided in mm9, we lifted the mouse full-stack annotation from mm10 to mm9. To do so, we first wrote the full-stack annotation in mm10 into a .bed file such that each line corresponds to a 200-bp segment. We then used the liftOver tool with default parameters to convert the 200-bp segments from mm10 to mm9. We filtered out regions in the lifted-over mm9 annotation that were mapped from >= 2 distinct segments in mm10.

For each full-stack state and each of the gene expression dataset (there are 38 of them with 2 replicates for each tissue type), we calculated the average gene expression of all genes that overlap with the state, while taking into account the genes’ length. We followed the same procedure described in Vu and Ernst, 2022. In particular, within a dataset, let the length and expression of gene *g* be denoted *L_g_* and *E_g_*, respectively. Let *B_s_* be the set of 200-bp genomic segments *i*’s that are assigned to state s in the mouse full-stack annotation, in mm9. Let *G_i_* denote the set of genes that overlap with genomic segment *i*. The gene-length-normalized average expression for state s is calculated as done previously [21]:

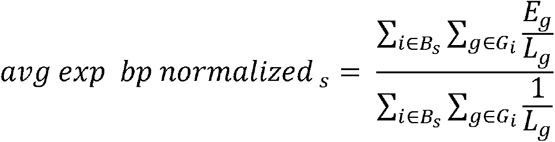

We then obtained the average gene expression for each full-stack state in each dataset. To calculate the average gene expression for the states in each of the 19 tissue types, we averaged the calculated average expression across the two replicate datasets for the same tissue type.

### External annotation sources

The sources for external annotations for enrichment analyses are as follows (all download links are listed in **Additional File 2**).

- Annotations of CpG islands, exon, gene bodies (exons and introns), transcription start (TSS), and transcription end sites (TES), 2kb windows surrounding TSSs (TSS2kb) in mm10 were RefSeq annotations included in ChromHMM (v 1.23) and originally based on annotations obtained from the UCSC genome browser [29,30] on July 26^th^, 2015.
- Annotation of coding gene regions correspond to coordinates of genes whose feature type is ‘CDS’ from GENCODE mm10 gene annotation, vM25 [31], accessed on February 3^rd^, 2022.
- Annotation of assembly gaps in mm10 were obtained from the UCSC genome browser and correspond to the Gap track [29,30], accessed on February 3^rd^, 2022.
- Annotations of pseudogenes in mm10 correspond to coordinates of genes whose gene type of transcript type contained ‘pseudogene’ from GENCODE’s mm10 gene annotation, vM25 [31].
- Blacklisted regions were downloaded from ENCODE project portal in mm10 from [32].
- Annotations of different repeat classes were downloaded from UCSC genome browser repeat masker track in mm10, accessed on Jan. 14^th^ 2022 [24].
- Annotations of Zinc finger genes in the mouse genome correspond to the coordinates of genes whose name contained ‘Zfp’ based on GENCODE mm10 annotation vM25 [31].
- Annotations of different chromosomes’ coordinates were downloaded from UCSC genome browser’s data of chromosome sizes in mm10, from https://hgdownload-test.gi.ucsc.edu/goldenPath/mm10/bigZips/mm10.chrom.sizes [29,30].
- LECIF scores measure that human-mouse conservation at functional genomics level, and were downloaded in version 1.1 from https://github.com/ernstlab/LECIF [25]. For each full-stack state, we reported the average LECIF score of overlapping genomic bases with the state
- CTCF peaks data were downloaded as bed files format from Mouse ENCODE Project [5,6]. We only included data files that has ‘*File analysis title*’ starting with ENCODE4 based on the metadata. In total, we obtained data of CTCF peaks for 42 ChIP-seq experiments from profiling CTCF in 28 unique biosamples. Details and download links for CTCF peaks data is available in **Additional File 2.**
- PhastCons conserved elements [26] based on the 60-way multi-species sequence alignment were downloaded from the UCSC genome browser (https://hgdownload.soe.ucsc.edu/goldenPath/mm10/database/phastConsElements60way.txt.gz).

## Supporting information

AF2_model_input_data

AF3_characterize_states

AF4_overlap_enrichments

AF5_relationship_with_ct_model

## Availability of data and materials

Mouse full-stack chromatin state annotation are available at https://github.com/ernstlab/mouse_fullStack_annotations in mm10. The code to analyze the fullstack states are available at https://github.com/ernstlab/mouse_fullStack_annotations. All download links for input for the mouse full-stack model are available in **Additional File 2**.

## Acknowledgements

We thank Soo Bin Kwon for helping us collect the data of per-cell-type chromatin state annotations. We thank members of the Ernst lab for helpful feedback in completing this manuscript.

## Ethics Declarations

Competing Interests: The authors declare that they have no competing interests.

## Supplementary Information

Additional File 1: Supplementary Figures S1-S9

Additional File 2: Metadata and download links for input data used for model learning, CTCF elements.

Additional File 3: Summary characterizations of mouse full-stack states.

Additional File 4: Mouse full-stack states’ average LECIF scores, enrichments with human fullstack states, repeat classes, chromosomes and CTCF elements.

Additional File 5: Mouse full-stack states’ relationships with per-cell-type annotations (supporting supplementary figures 6-7)

