## Supplementary material for "Universal chromatin state annotation of the mouse genome": AF1_supp_figures

**List of supplementary figures**

**Supplementary Figure 1:** Mouse full-stack states transition probabilities.

**Supplementary Figure 2:** Positional enrichments of full-stack states around annotated transcription start sites and transcription end sites.

**Supplementary Figure 3:** Mouse full-stack states enrichments with different chromosomes. *(An excel sheet supporting this figure is in Additional File 4).*

**Supplementary Figure 4:** Mouse full-stack states enrichments with different classes of repeats. *(An excel sheet supporting this figure is in Additional File 4).*

**Supplementary Figure 5:** Enrichment of select mouse full-stack states with different classes of repeat elements.

**Supplementary Figure 6:** Full-stack states maximum-enrichments with annotated concatenated-model chromatin states in 66 mouse reference epigenomes. *(An excel sheet supporting this figure is in Additional File 5).*

**Supplementary Figure 7:** Estimated probabilities of per-cell-type concatenated-model chromatin states overlapping with mouse full-stack states. *(An excel sheet supporting this figure is in Additional File 5).*

**Supplementary Figure 8:** Enrichments of mouse full-stack states with human full-stack states. *(An excel sheet supporting this figure is in Additional File 4).*

**Supplementary Figure 9:** Mouse full-stack states’ relationship with LECIF scores, human full-stack states and phastCons elements. *(An excel sheet supporting this figure is in Additional File 4).*

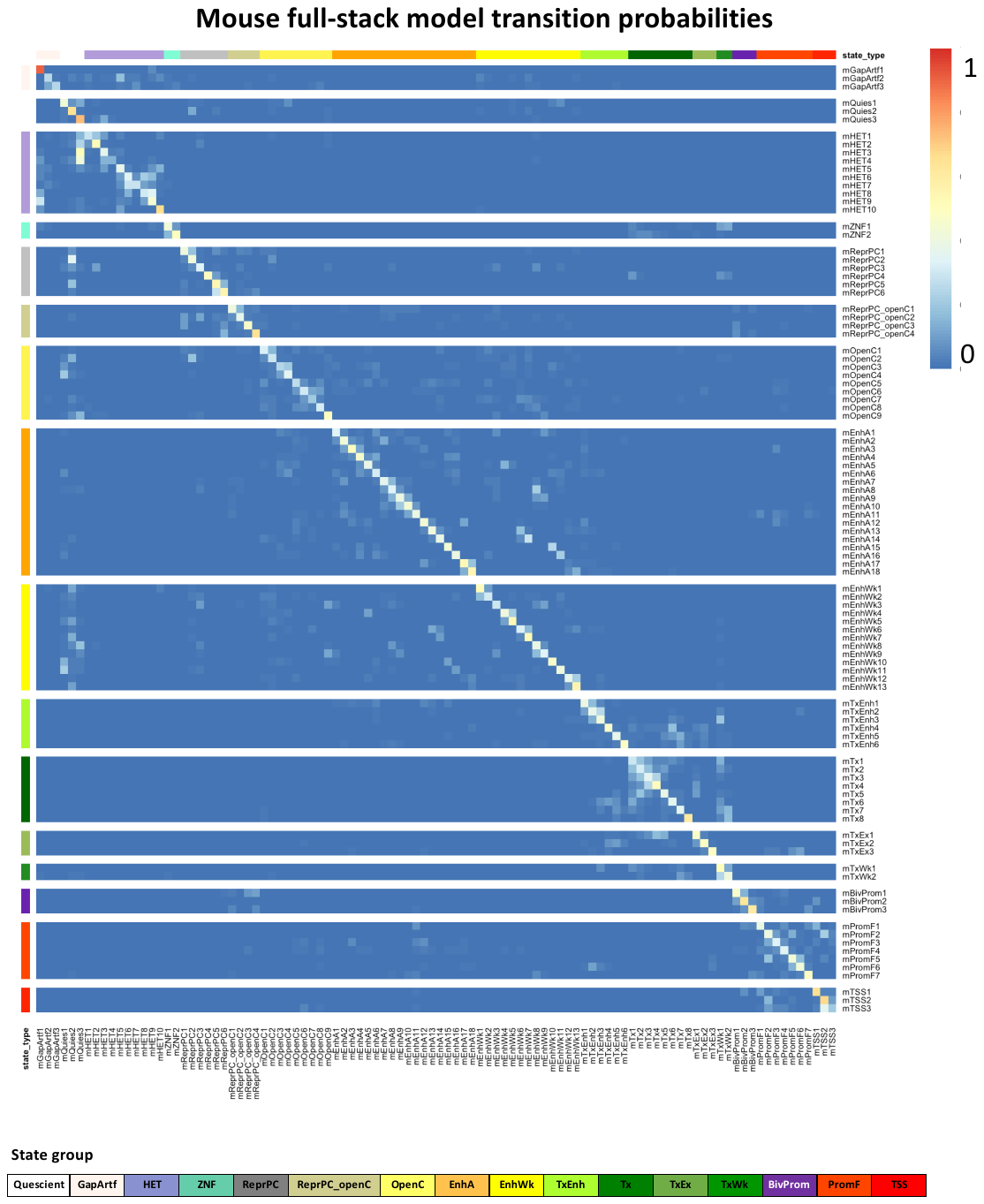

**Supplementary Figure 1: Mouse full-stack states transition probabilities.** Each row and each column correspond to a full-stack state, ordered based on their associated state group. The heatmap shows for each state assigned at a current genomic position (rows) the probabilities of transitioning to another state (columns) at the subsequent genomic position. The state groups are shown at the bottom.

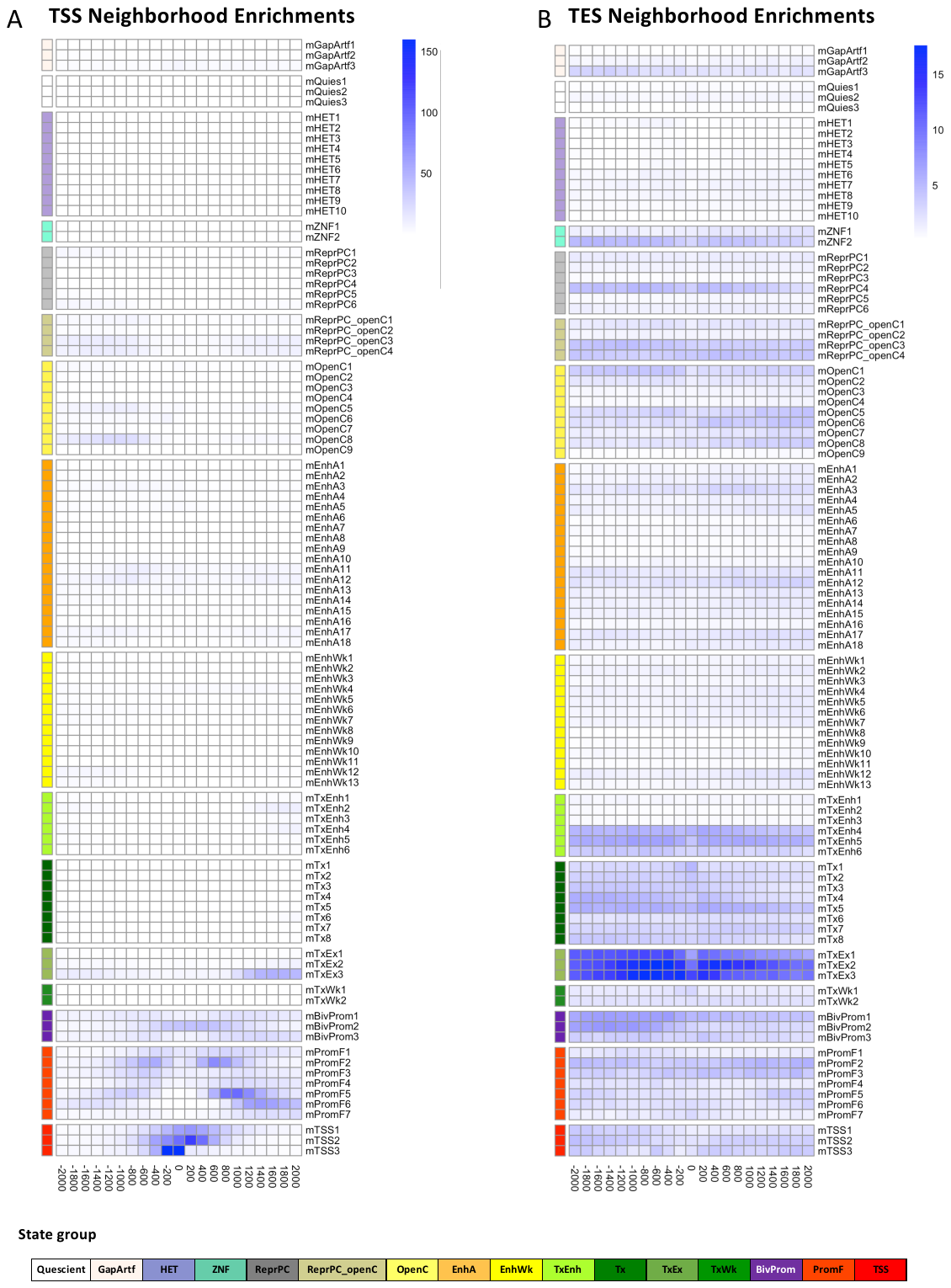

**Supplementary Figure 2: Positional enrichments of full-stack states around annotated transcription start sites and transcription end sites.** The figure shows positional fold enrichments for positions within 2kb of annotated (**a**) transcription start sites (TSS) and (**b**) transcription end sites (TES). Each column corresponds to one 200bp window as indicated at bottom. Positive coordinate values represent the number of bases downstream in the 5’ to 3’ direction of transcription, while negative values represent the number of bases upstream. Enrichments are calculated based on a genome-wide background. Color scale of enrichments is indicated at right for each panel. State groups’ color legends are shown at the bottom.

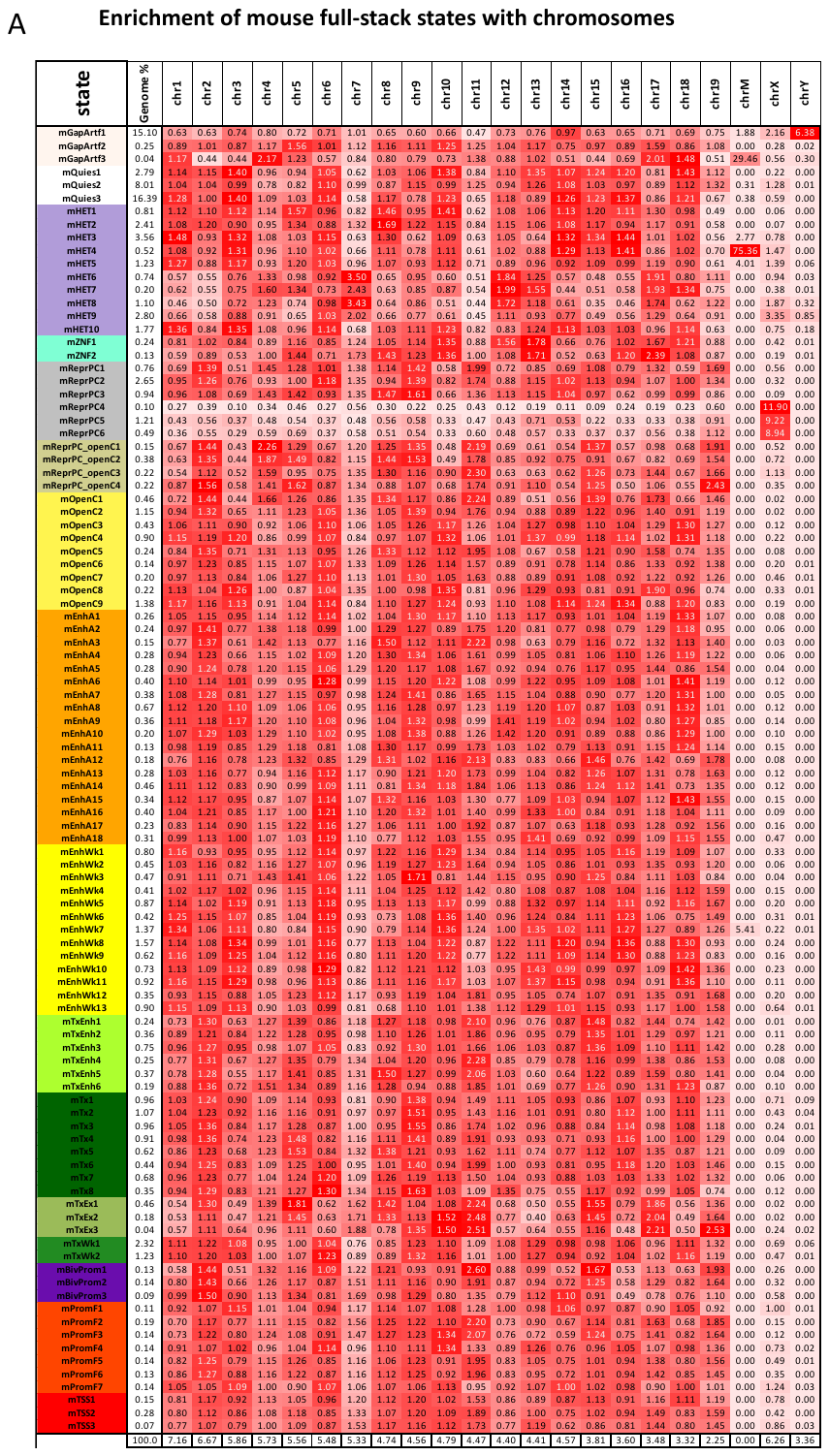

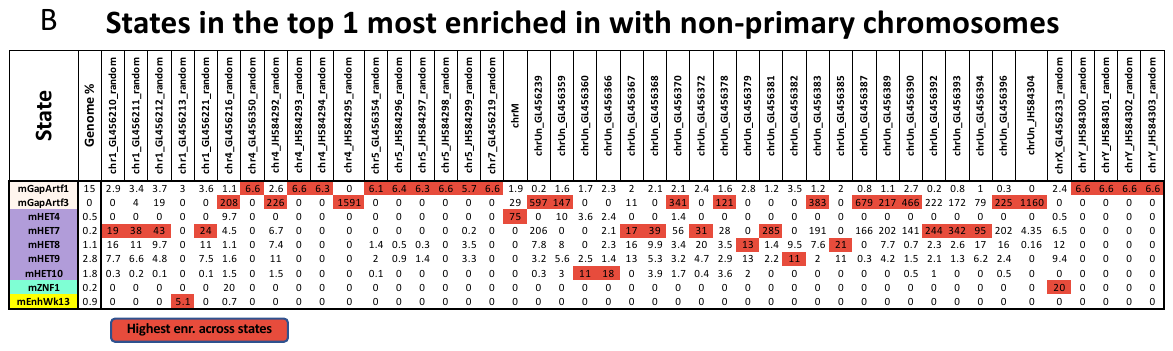

**Supplementary Figure 3: Mouse full-stack states enrichments with different chromosomes. (A)** The first and second columns show the mouse full-stack states and their percent genome coverage. The following columns correspond to different chromosomes. The heatmap shows fold enrichments of each state with each chromosome. Coloring of the heatmap is column-specific. The last row shows the percentage of the genome that each chromosome covers. Certain states in polycomb repressed group (mReprPC4-6) show distinctly high enrichments with chromosome X. **(B)** The first and second columns show mouse full-stack states and their genome coverage, respectively. The following columns correspond to different scaffold chromosomes. Only states that show highest enrichments with at least one scaffold chromosome are shown. Within each column, the highest enrichment values across 100 mouse full-stack states are colored red. States in ‘assembly gaps and alignment artifacts’ or in heterochromatin groups show highest enrichments with multiple scaffold chromosomes.

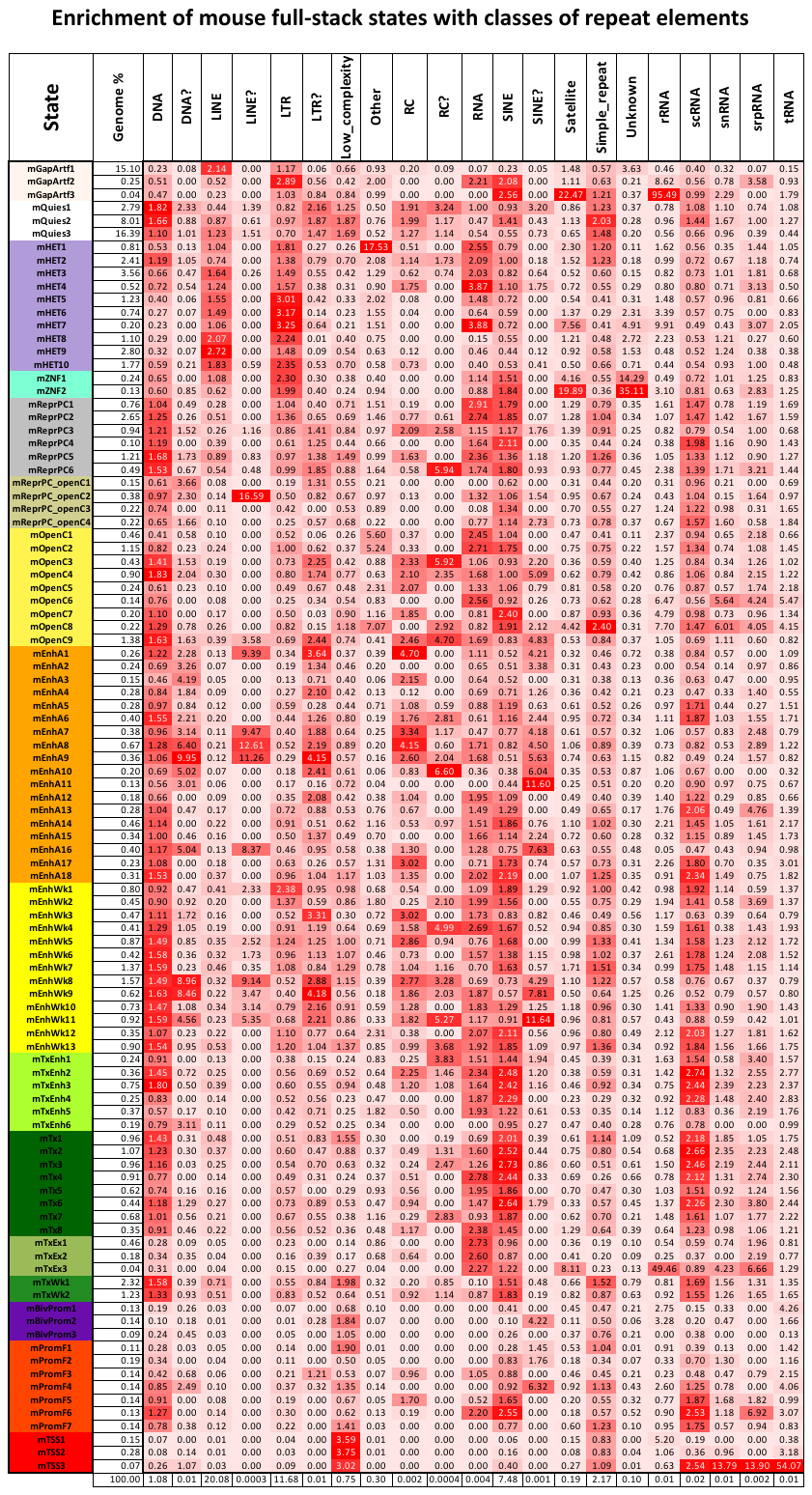

**Supplementary Figure 4:** **Mouse full-stack states enrichments with different classes of repeats** (Smit *et al.*, 2015). The first and second columns show the mouse full-stack states and their genome coverage. The following columns correspond to different repeat classes. The following columns correspond to different classes of repeat elements (with elements named with ‘?’ excluded). The heatmap shows fold enrichments of each state with each repeat class. Coloring of the heatmap is column-specific. The last row shows the percentage of the genome that each repeat class covers.

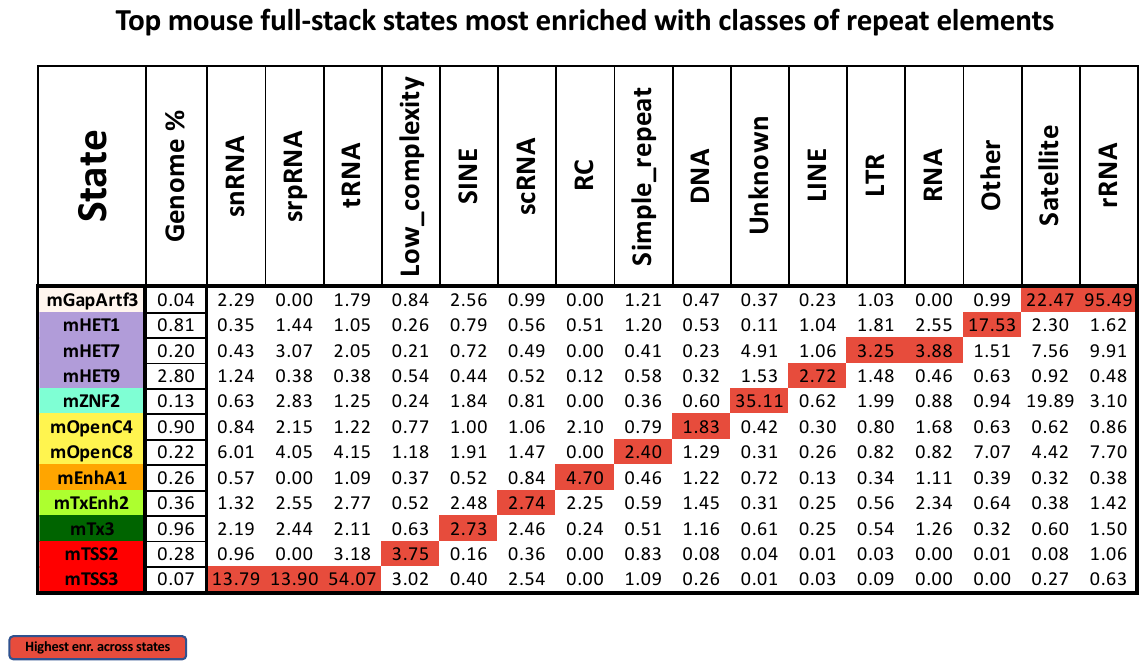

**Supplementary Figure 5: Enrichment of select mouse full-stack states with different classes of repeat elements** (Smit *et al.*, 2015). The first and second columns show mouse full-stack states and their genome coverage, respectively. The following columns correspond to different classes of select repeat elements. *Only states that show highest enrichments with at least one repeat class are shown*. Within each column, the highest enrichment values across 100 mouse full-stack states are colored red.

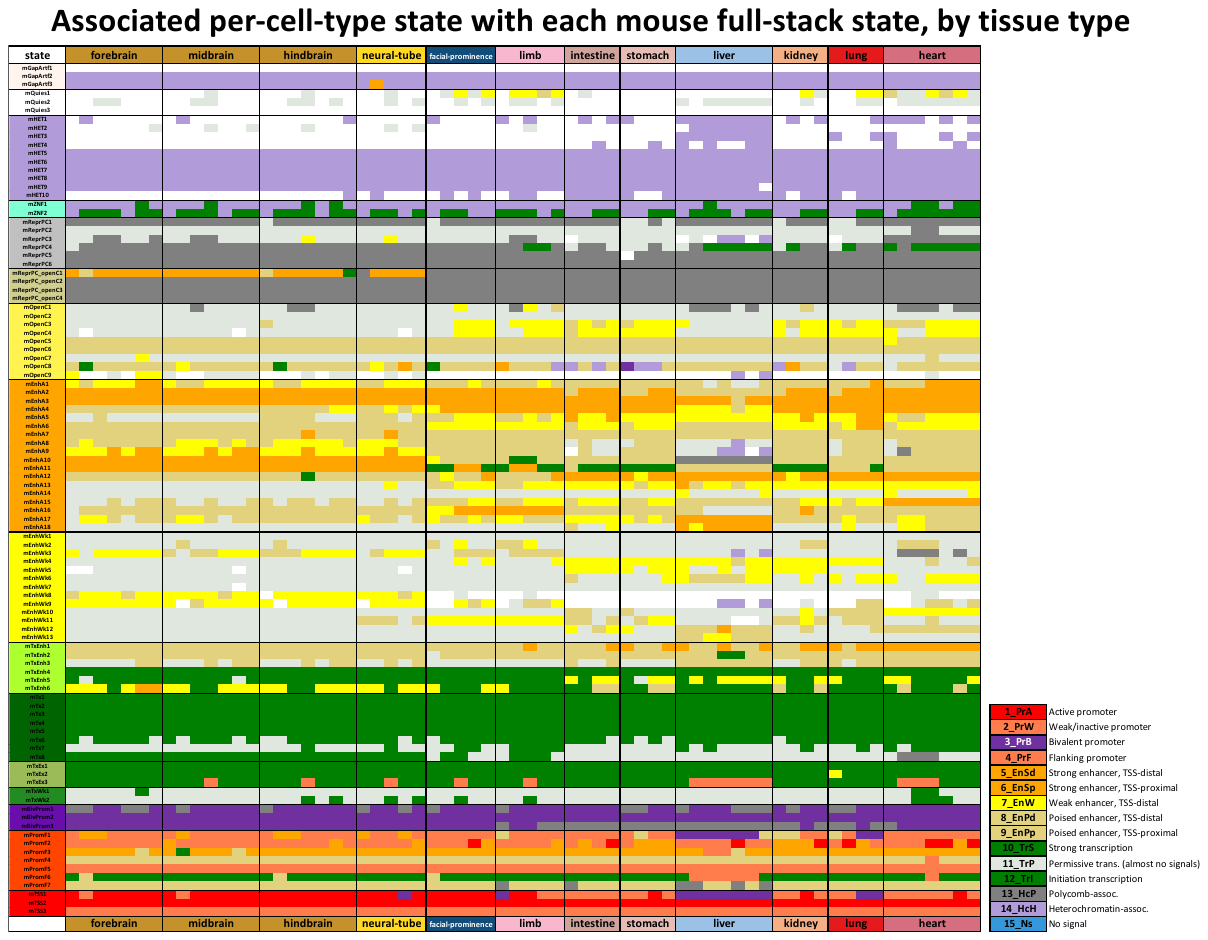

**Supplementary Figure 6: Full-stack states maximum-enrichments with annotated concatenated-model chromatin states in 66 mouse reference epigenomes** (Gorkin *et al.*, 2020)**.** Each row corresponds to one of 100 mouse full-stack state (**Methods**). Each column corresponds to a reference epigenome, grouped by the associated cell types as colored at the top and bottom. Each color entry corresponds to a reference epigenome and mouse full-stack state combination. The color corresponds to the chromatin state from the concatenated 15-state model annotating the respective mouse reference epigenome that is most enriched with the respective mouse full-stack state. Description of states in the per-cell-type 15-state concatenated model is in the bottom (Gorkin et al., 2020). The figure highlights how some mouse full-stack states are maximally enriched with the same concatenated-model chromatin states across all the reference epigenomes; for example, states mTx1-5 are maximally enriched with the strong transcription state in all 66 reference epigenomes’ 15-state concatenated annotation. Other mouse full-stack states are enriched for distinct concatenated states in different cell types, for example state mEnhA17-- characterized as an enhancer state in liver, spleen and bone marrow based on emission probabilities of enhancer associated marks-- is most enriched with an active enhancer in liver cell types, while being most enriched with poised/weak enhancer states in others. Detailed description of each mouse full-stack state enrichment patterns with concatenated states can be found in **Additional File 5**.

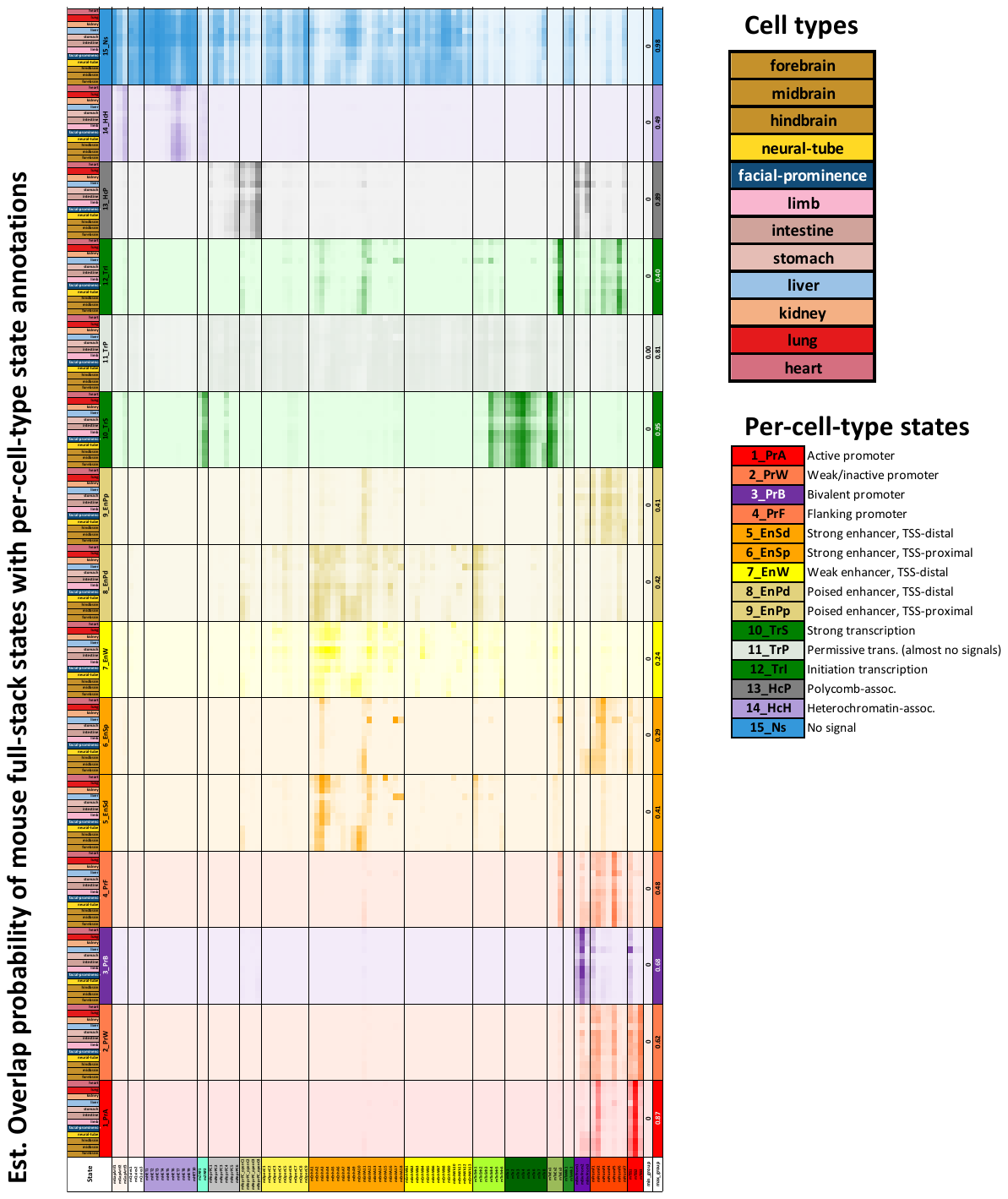

**Supplementary Figure 7: Estimated probabilities of per-cell-type concatenated-model** **chromatin states overlapping with mouse full-stack states.** The figure shows estimated probabilities of per-cell-type chromatin state annotations overlapping with mouse full-stack states observed in different cell groups (Gorkin *et al.*, 2020). This figure is also provided as an excel file in **Additional File 5**. The figure is based on a 15-state per-cell type chromatin state model trained on 66 mouse reference epigenomes from 12 cell groups (Gorkin *et al.*, 2020). Each row corresponds to a combination of per-cell type state (among 15 states) and cell group, as denoted in the first two columns and legends on the right and matching with the colors in **Fig. S5**. We note that we changed here the concatenated-model no-signal state from white to blue for better visibility. Rows corresponding to the same per-cell-type model state are grouped together (into 15 bigger rows). The 100 following columns correspond to 100 mouse full-stack states. Values in the heatmap correspond to the estimated probability a genomic position annotated as a mouse full-stack state (column) is also annotated as a concatenated-model state in a reference epigenome from the corresponding cell group (row) (**Methods**). The last two columns show the minimum and maximum probabilities observed for each per-cell type state for any combination of tissue group and mouse full-stack state. The heatmap colors correspond to the 15-state’s colors and are scaled such that the maximum probability value in each row block is colored darkest (as seen in the right most column). The figure complements **Fig. S5** in providing information on how each full-stack state can correspond to different per-cell-type states, hence stratifying mouse full-stack states’ characteristics in more details. For example, mouse full-stack state mTSS1 shows high probabilities of overlapping bivalent promoter state in liver cells, and moderate probabilities of overlapping the flanking/weak promoter state in other cell groups.

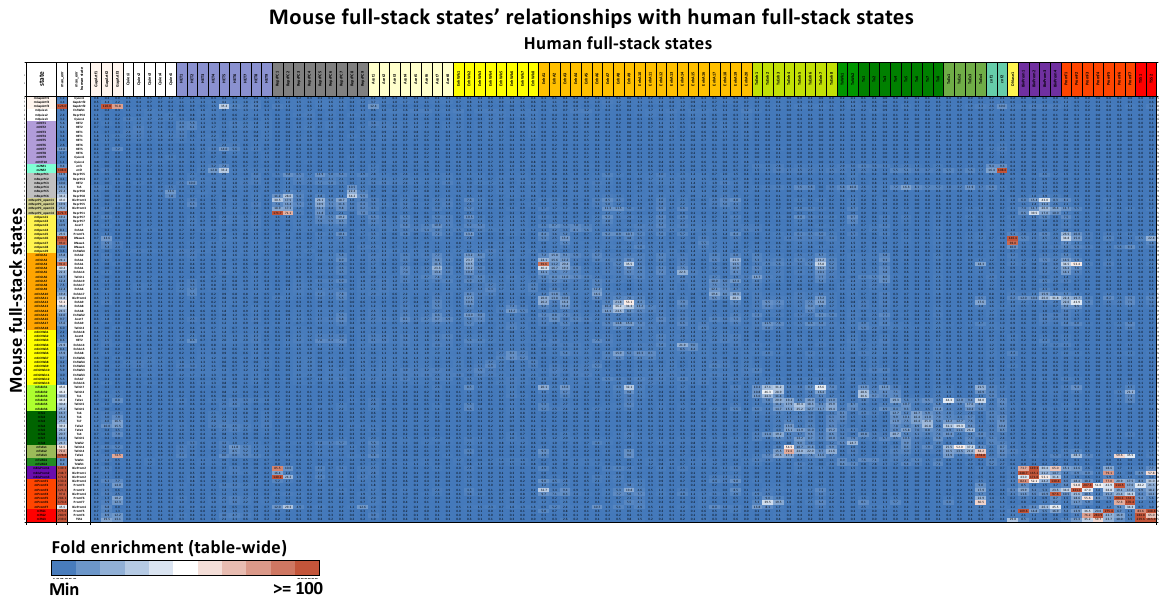

**Supplementary Figure 8: Enrichments of mouse full-stack states with human full-stack states (Vu and Ernst, 2022).** The first three columns show mouse full-stack states, the maximum fold enrichment with a human full-stack state (across all human states) and the corresponding human state, respectively. The following columns show the overlap enrichments with of each mouse state (rows) with each human state (columns). Across all pairs of states, the smallest enrichment values are colored blue and enrichment values >= 100 are colored red. This figure is also provided in **Additional File 4.**

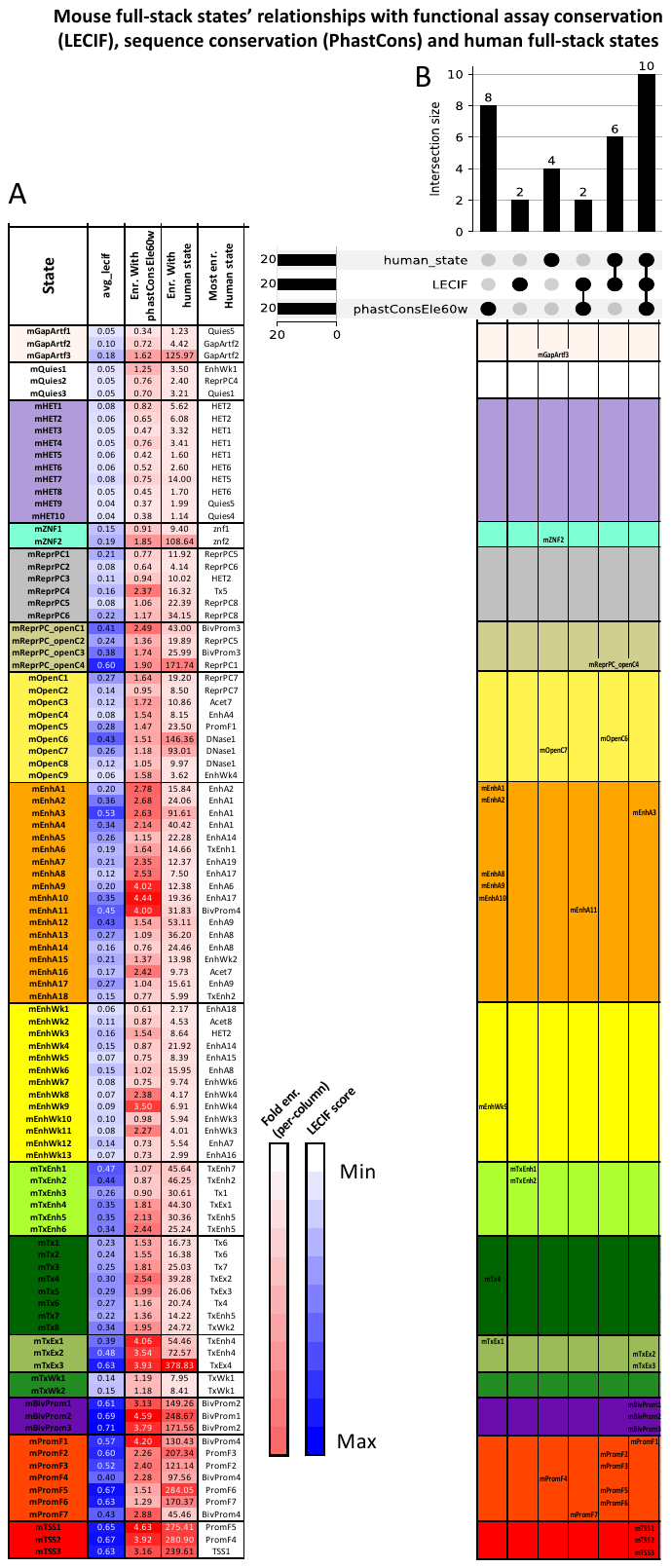

**Supplementary Figure 9: Mouse full-stack states’ relationship with LECIF scores, human full-stack states and phastCons elements.** LECIF scores were developed to measure the level of evidence of human-mouse conservation at functional/epigenomic levels, with higher score (maximum of 1 and minimum of 0) implies higher evidence of conservation (Kwon and Ernst, 2021). PhastCons elements correspond to genomic regions showing strong 60-way multi-species sequence alignment conservation (Siepel *et al.*, 2005). Human full-stack states were learned from >1,000 Chip-seq/DNase-seq datasets in human, and provide annotation of the human genome that is shared across cell/tissue types (Vu and Ernst, 2022). **(A)** The heatmap shows mouse full-stack states (rows)’ average LECIF scores, enrichments with phastCons elements and the maximum enrichments with human full-stack states. The first and second columns show the mouse full-stack states, and the percentage of the genome that each state covers. Coloring of the next 3 columns is column specific, as specified in legend. The last row shows the percentage of the genome that each LECIF score range covers. **(B)** Upset plot showing the number of states that are among the top 20 states with either (1) highest average LECIF score, or (2) hig`hest enrichments with PhastCons elements or (3) highest maximal enrichments with human full-stack states. Within each category, the column below the upset plot lists states that are in the top 20 most associated (as measured by average LECIF scores or fold enrichments) with the combination enrichment contexts.
